# Comparison of classification methods for tissue outcome after ischemic stroke

**DOI:** 10.1101/551903

**Authors:** Ceren Tozlu, Brice Ozenne, Tae-Hee Cho, Norbert Nighoghossian, Irene Klærke Mikkelsen, Laurent Derex, Marc Hermier, Salvador Pedraza, Jens Fiehler, Leif Østergaard, Yves Berthezène, Jean-Claude Baron, Delphine Maucort-Boulch

## Abstract

In acute ischemic stroke, identifying brain tissue at high risk of infarction is important for clinical decision-making. This tissue may be identified with suitable classification methods from magnetic resonance imaging (MRI) data. The aim of the present study was to assess comparatively the performance of five popular classification methods (Adaptive Boosting (ADA), Logistic Regression (LR), Artificial Neural Networks (ANN), Random Forest (RF), and Support Vector Machine (SVM)) in identifying tissue at high risk of infarction on human voxel-based brain imaging data. The classification methods were used with eight MRI parameters including diffusion-weighted imaging (DWI) and perfusion-weighted imaging (PWI) obtained in 55 patients. Sensitivity, specificity, the area under the receiver operating curve (ROC) as well as the area under the precision-recall curve criteria were used to compare the method performances. The methods performed equally in terms of sensitivity and specificity while the results of the area under the ROC were significantly better for ADA, LR, ANN and RF. However, there was no statistically significant difference between the performances of these five classification methods regarding the area under the precision-recall curve, which was the main comparison metric.

## 1. INTRODUCTION

Ischemic stroke is one of the major causes of death or long-term disability in most developed countries.^1^ One of the current medical stroke therapies is intravenous thrombolysis which has to be administered as soon as possible after symptom onset. Besides, identifying the tissue at risk of infarction with an imaging tool would help decision-making in personalized treatment.^2^

Brain imaging based on Magnetic Resonance Imaging (MRI) provides important variable related to acute cerebral ischemia. Especially, diffusion-weighted imaging (DWI) is a marker of irreversibly damaged tissue and perfusion-weighted imaging (PWI) presents regions of reduced tissue blood supply and particularly the ischemic penumbra, which is at risk of infarction but still salvageable by early reperfusion.^3,4^ In the acute stroke setting, a marked discrepancy between the size of abnormal tissue identified by DWI vs. PWI^1^, so called DWI-PWI mismatch, is often encountered, and the performance of DWI and PWI data combined in identifying the tissue at risk of infarction has been found to be superior to that of DWI data or PWI data alone.^3,4^

Identifying the tissue at risk of infarction in each patient using MRI and specific statistical methods would help to determine the subject’s most appropriate treatment.^3,4^ Many classification methods have been already used to provide the risk of infarction on voxel-based data. For example, generalized linear models have been proposed to estimate the probability of infarction on the basis of diffusion- and perfusion-weighted images from humans^5,6^, but machine learning algorithms may outperform the generalized linear model in case of complex multimodal data. Artificial Neural Networks (ANN) was used on animal and human imaging data and reported promising findings regarding predicting the outcome of ischemic tissue^7,8^. In 2011, another study from the same authors^9^ found better results with Support Vector Machine (SVM) than with ANN on animal data. Random Forest (RF) was also used by many studies^10–12^ for the prediction of the infarction from imaging data and Livne et al.^10^ showed that boosted tree models gave higher performance compared to generalized linear models. An extensive study by Bouts et al.^8^ compared five classification methods on experimental animal data; the five methods gave similar results. However, extrapolating these findings to man may not be appropriate.^9^ Thus, Winder et al.^15^ compared the prediction accuracy of three classification methods (nearest-neighbor, generalized linear model, and RF) using human voxel-based stroke data; RF --as a machine learning algorithm--performed significantly better than the two other methods.

The aim of the present study was to assess the performance of five highly used classification methods in the literature (Adaptative Boosting – ADA, ANN, Logistic Regression – LR, RF, and SVM) in identifying the tissue at high risk of infarction from voxel-based human data.

## 2. MATERIAL AND METHODS

### 2.1. Patients

The study used data of patients from the I-KNOW multicentre study that included prospectively patients who underwent MRI at admission and follow-up to estimate voxel-based probabilistic maps of infarction risk.^11^ Patients with lacunar or posterior circulation stroke, unknown time of stroke onset, unknown T2 FLAIR sequence measure or intracerebral haemorrhage on MRI were excluded. Overall, 55 patients were used for each classification method. The inclusion criteria were: (1) National Institutes of Health Stroke Scale (NIHSS) ≥4; (2) DWI and PWI data consistent with an anterior-circulation acute ischemic stroke; (3) admission MRI carried out within 6 hours in case of intravenous recombinant tissue plasminogen activator (rt-PA) use or within 12 hours in case of conservative treatment. The I-Know study conformed with the Helsinki Declaration, the rules laid out by the Council of Europe Convention on Human rights and Biomedicine, Directive 95/46/EC of the European Parliament and of the Council of 24 October 1995 on the protection of individuals with regard to the processing of personal data and on the free movement of such data, and with the legislation and regulations in Denmark, Germany, France, and Spain, respectively. The study was approved by the Aarhus, Hamburg, Lyon, and Girona hospitals respective regional ethics committees, and carried out after informed consent from the patients.

### 2.2. Image acquisition and processing

On admission, all patients underwent: i) DWI MRI (3 or 12 directions, repetition time >6000 ms, field of view 24 cm, matrix 128×128, slice thickness 3 or 5mm); ii) a Fluid-Attenuated-Inversion-Recovery (FLAIR: repetition time 8690 ms, echo time 109 ms, inversion time 2500 ms, flip angle 150°, field of view 21 cm, matrix 224×256, 24 sections, section thickness 5 mm, slice gap 1 mm); iii) a PWI MRI (echo time 30-50 ms, repetition time 1500ms, field of view 24 cm, matrix 128×128, 18 slices, thickness 5 mm, gap=1 mm, gadolinium-based contrast at 0.1 mmol/kg, intravenous injection 5 mL/s followed by 30 mL saline).

Diffusion-weighted sequence generated maps of DWI and apparent diffusion coefficient (ADC) parameters. PWI tracked the bolus of injected contrast agent to generate maps of cerebral blood volume (CBV), cerebral blood flow (CBF), and temporal parameters such as the time to peak (TTP), the mean transit time (MTT), and the time-to-maximum (TMAX). Perfusion maps were computed by circular singular value decomposition of the tissue concentration curves with an arterial input function from the contralateral middle cerebral artery. Using a reference region from the contralateral normal white matter, temporal parameters normalized by subtracting the mean contralateral value and all further references to MTT, TMAX, and TTP refer to the relative parameters.

The parameter maps of each patient were normalized using the mean and the standard deviation of the contralateral tissue (cerebrospinal fluid excluded). The means were calculated on three consecutive slices and the standard deviation on the full brain volume. The parameter values were then centred and scaled to be comparable in effect size.

All eight MRI-based parameters (i.e., T2 FLAIR, ADC, CBV, CBF, TTP, MTT, TMAX, and DWI) were used as predictors to characterize each voxel’s risk of infarction. FLAIR imaging at one month after stroke onset was used to contour the boundaries of the irreversibly damaged brain tissue in stroke patients by a clinical. FLAIR at one month was used as outcome to compare the predictions of the methods.

### 2.3. Classification methods

Four machine learning methods (SVM, ANN, RF, and ADA) and LR were used in this study. (For more details of methods and their settings, see the Appendix). These five methods allow identifying the risk of infarction of each voxel-based observation.

The LR estimated the infarction risk of a voxel as its probability of being infarcted after certain follow-up periods using the same above-cited combination of eight MRI parameters.^12^

The SVM separated the observations into healthy or infarcted using a linear border. The closest observations to the border in each class were named support vectors. Then, the support vectors helped to choose the best border either through maximizing the distance between the border and the support vectors or through minimizing the number of misclassified observations.^13^ The signed distance to the border is related to the risk of infarction. A zero distance corresponds to an observation located on the border, a positive distance to a high risk, and a negative distance to a low risk.

The ANN constitute a mathematical representation of natural neural networks.^14^ Each network is composed of several types of layers: (1) an input layer composed of all data, (2) an output layer giving the final outcome, and (3) one (or more) hidden layers between the input and the output layer that consist(s) of a set of neurons that process the data and are connected to the input and output layers. The input layer sends first the information to the next layer with an initial weight and the weight is updated after the response of the network is reached in the output layer. The update iterations continue until there is little change. The weighted sum of the responses of the neurons in the output layer is then computed. The risk of infarction is found by applying a sigmoid function to this weighted sum.

The other classification methods, RF and ADA, are ensemble methods that combine several decision trees. RF builds several bootstrap samples from the original data and fits a decision tree to each sample to classify the observations into healthy or infarcted.^15,16^ At each observation, the risk of infarction is then computed as the percentage of trees that classify this observation as infarcted. The ADA weights the misclassified observations using a set of decision trees.^16^ The classification of the first tree is performed with the same weight for each observation but the weights of the misclassified observations are increased after each decision tree classification. The performance of each tree is computed using the misclassification error of the decision tree and the weights given to the observations. At each observation, the infarction risk is the result of the classification of each tree weighted by its performance.

### 2.4. Statistical analysis

The performance of the five classification methods tested here were mainly compared in terms of area under the precision-recall curve *(AUC_prc_*). This criterion was selected because it summarizes the predictive ability of each method over all possible thresholds allowing for infarction prevalence. The *AUC_pr_* was preferred to the area under the receiver operating curve *(AUC_roc_)* because the ischemic stroke data were highly imbalanced; i.e., the infarcted voxels are much less numerous than non-infarcted voxels.^17^ However, as many studies have already used the *AUC_roc_* to compare classification methods, this criterion was also reported here to allow result comparisons.

The leave-one-out cross-validation approach was used to reduce overfitting and provide an accurate estimation of the prediction performances of the different methods.^18^ Each model was trained with two loops (outer and inner) of leave-one-out cross validation (k = 55 for both loops). The outer loop (repeated over 100 random partitions of the full dataset) provided a training set for model building and test set for model assessment. The inner loop (repeated over 55 different partitions of the training dataset only) performed grid-search to find the set of model hyperparameters that minimized the misclassification rate. Afterwards, the prediction relative to the removed patient was performed using the same method. *AUC_prc_,AUC_roc_*, sensitivity, and specificity were separately calculated for each of the 55 patients and the medians of these values given as evaluation criteria.

The results from the different methods enabled classifying each voxel into healthy or infarcted using a given threshold. The threshold was defined as the value that minimizes the difference between the observed and the predicted infarcted volume; thus, a voxel with an infarction risk above the threshold was considered as infarcted. Threshold was calculated in leave-one-out cross validation for each outer loop to avoid overfitting. The predicted infarction volume for each patient could be calculated as the sum of the voxels classified as infarcted.

Method performances calculated with *AUC_pr_, AUC_roc_*, sensitivity, and specificity were compared with the Kruskal-Wallis test. An higher value of *AUC_pr_, AUC_roc_*, sensitivity, and specificity obtained by a classification method indicates a better prediction performance. The independence between treatment and gender, having hypertension, and cigarette smoking before the stroke was tested with Chi-Squared test. The age between treated and untreated patients was compared with unpaired two-samples Wilcoxon rank sum and signed rank test. The significance level set at p < 0.05. The free software R (https://www.r-project.org) with 3.4.4. version was used for all statistical analyses and graphs.

## 3. RESULTS

### 3.1. Patients

Table 1 shows the patients’ characteristics. We observed that 37 out of the 55 patients were treated with intravenous thrombolysis. The number of female and male was similar among treated patients (49% vs 51%) whereas there were more men than female among untreated patients (22% vs 78%). The majority of patients (65%) had hypertension before the stroke. The gender and receiving treatment were significantly independent (Chi-Squared test, p-value=0.11). 30% of the patients had smoked regularly before stroke. More untreated patient has been observed smoking than treated patient (50% vs 21%) but there was no significant relationship between smoking and the treatment (p-value=0.25). In terms of infarcted volume, the median of final infarction volume was higher than the DWI lesion volume. However, we observed no significant change in infarcted volume at one month for treated and untreated patients (Paired Wilcoxon test: *p* — *value_treated_* = 0.08, *p* — *value_untreated_* = 0.26).

**Table 1.** Patients’ clinical characteristics at hospital admission and one month later. The patients are divided into two groups such as treated with rt-PA or untreated with rt-PA Values are presented as median [first quartile; third quartile] – rt-PA: recombinant tissue plasminogen activator.

|  | Treated<br>(n=37) | Untreated<br>(n=18) | All patients<br>(n=55) |
| --- | --- | --- | --- |
| Female | 49% | 22% | 40% |
| Prior hypertension | 62% | 72% | 65% |
| Smoking | 21% | 50% | 30% |
| Age (yrs) | 70 [65; 77] | 67.5 [60; 74,75] | 68 [62; 77] |
| DWI lesion volume, mL | 12 [4; 27] | 4 [3; 21] | 11 [4; 26] |
| Final infarct volume, mL | 12 [3; 43] | 8 [2, 24] | 10 [3; 40] |

### 3.2. Classification

Table 2 presents the *AUC_prc_, AUC_roc_*, sensitivity, and specificity values relative to the five classification methods. The *AUC_prc_* values ranged between 0.20 and 0.34; the highest value was obtained with SVM and the lowest with LR. However, there was no statistically significant difference between *AUC_prc_* values (p-value = 0.75). All *AUC_roc_* values were higher than 0.7. RF, ADA, ANN, and LR performed the same and significantly better than SVM when *AUC_roc_* values compared with Kruskal-Wallis test (p-value <0.05). The sensitivity values were all less than 0.5 whereas the specificity values were all larger than 0.9. There was no statistically significant difference between methods in terms of sensitivity or specificity.

**Table 2.**
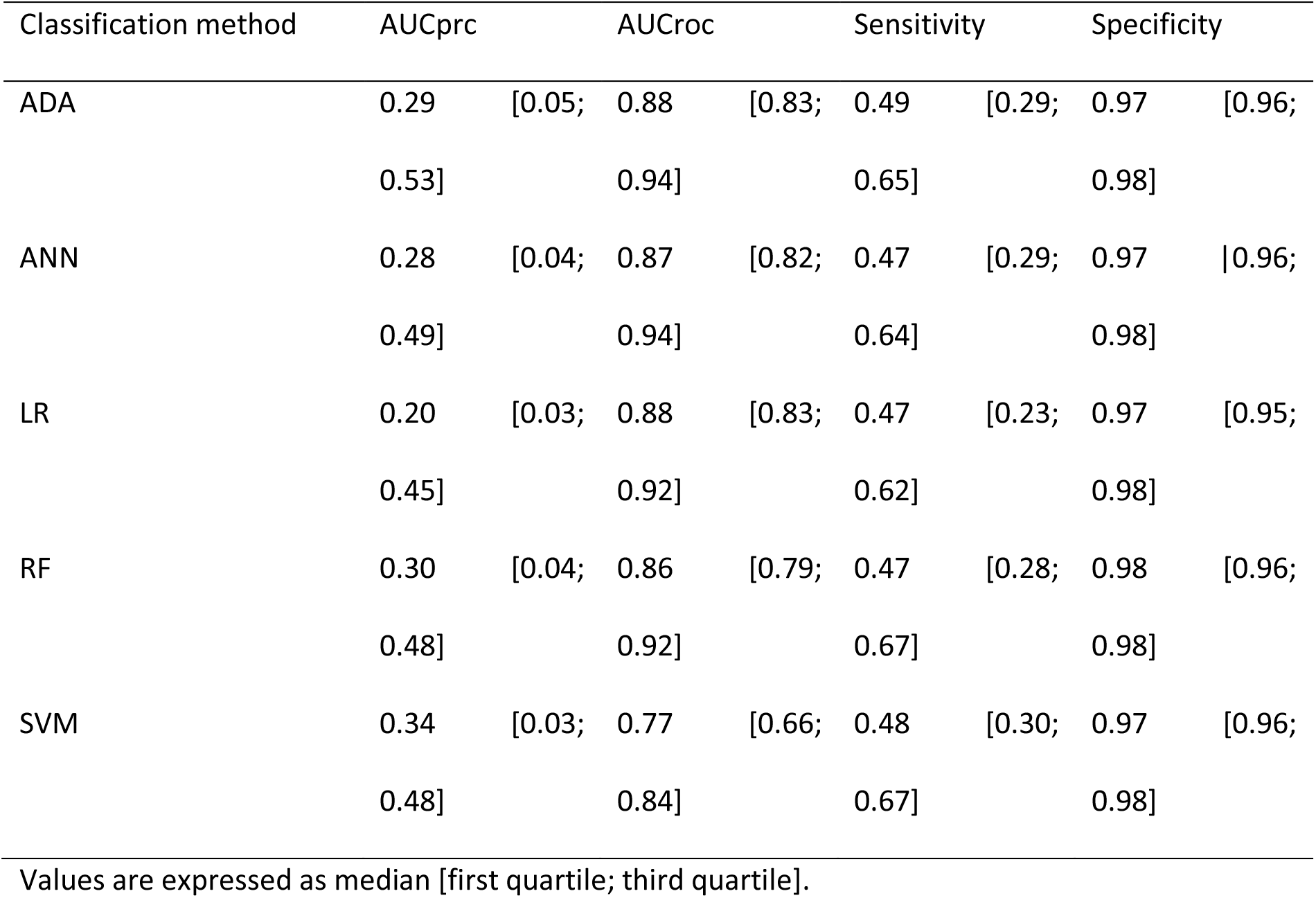
Performance criteria of the five studied classification methods.

Figure 1 shows the *AUC_prc_* and *AUC_roc_* values performed by LR and SVM on each patient data. These two methods were selected because they gave respectively the highest and lowest *AUC_prc_* values (Table 2). The graphs show that both *AUC_prc_* and *AUC_roc_* values with LR are higher than SVM values on almost all patient data.

**Figure 1.**
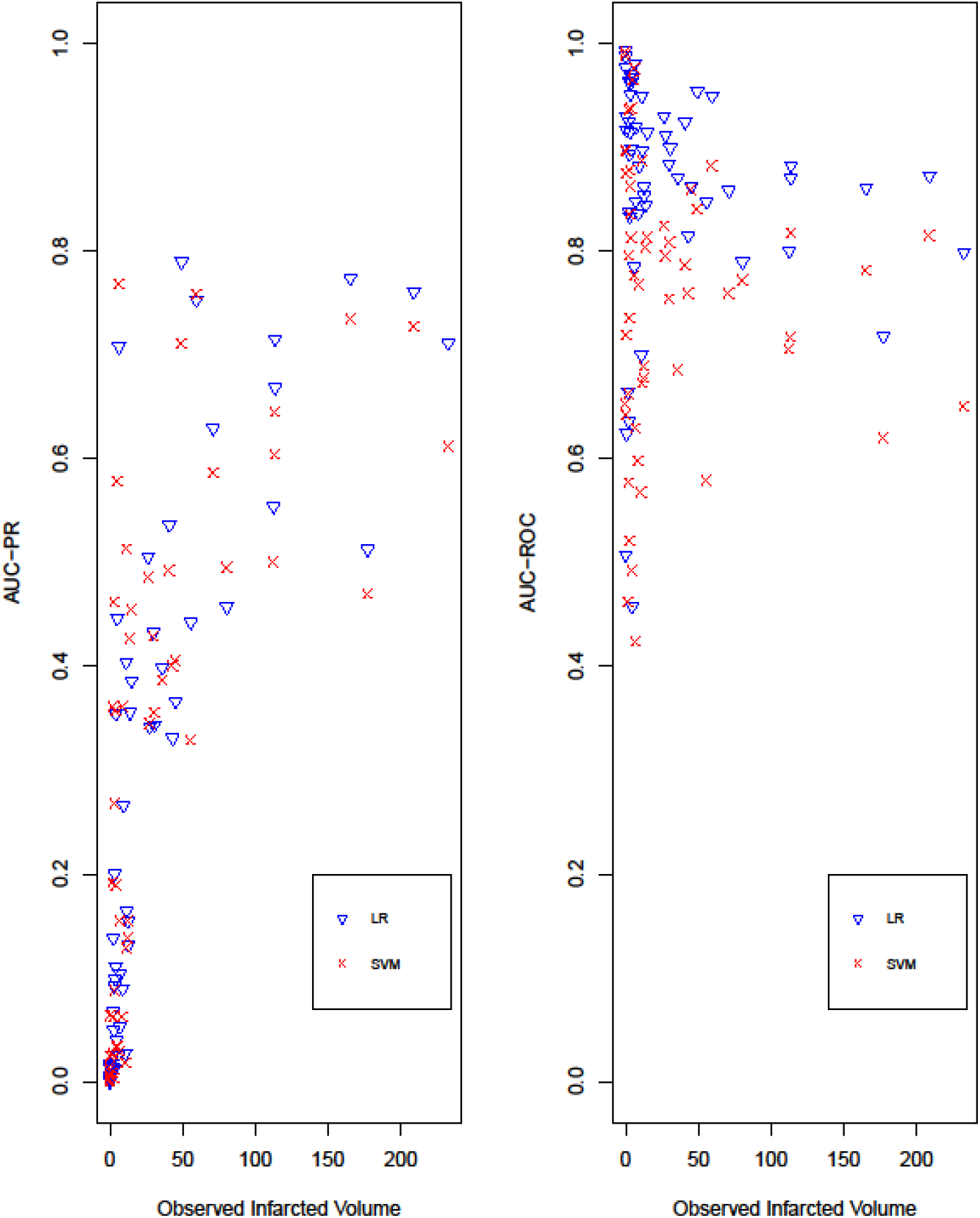
Observed infarcted volumes in 55 patients at one month after stroke as expressed by the area under precision-recall curve *AUC_prc_* (left panel) and the area under the receiver operating curve *AUC_roc_* (right panel) obtained with logistic regression (LR) and Support Vector Machine (SVM).

In the case of volumes <50 mL, both *AUC_prc_* and *AUC_roc_* values were highly variable whereas *AUC_roc_* values tended to have higher values than *AUC_prc_* values. In volumes > 100 mL, *AUC_prc_* and *AUC_roc_* values were similar.

Figure 2 shows the voxels of predicted infarction on illustrative examples of brain MRI slices. The rows show findings for two representative patients with respectively regularly and irregularly-shaped infarction. ADA, ANN, LR, and RF performed approximately the same in the two patients. For the patient shown in the top row, ADA, ANN, LR, and RF overestimated infarct size, with numerous false positive voxels. In the bottom row patient, all methods, particularly SVM, underestimated final infarct size.

**Figure 2.**
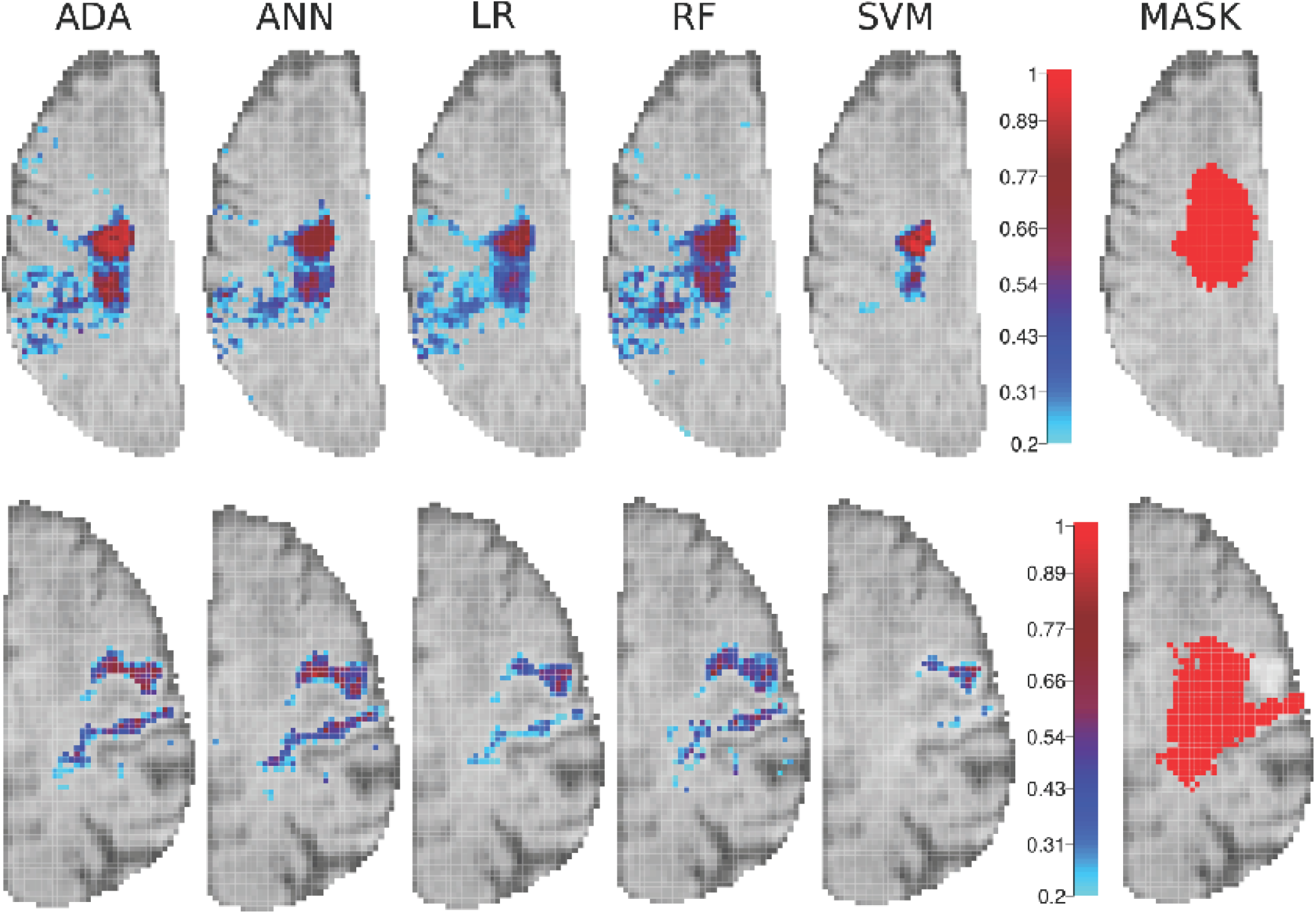
Voxel-wise infarction risk prediction ([0; 1] probability) obtained with the five methods (left), and final infarct (red) as delineated on one-month post-stroke FLAIR on the right hand side, for two representative patients. Each column shows the results yielded by each method, projected onto the template MRI mask. The risk scale ranges from light blue (risk = 0) to deep red (risk = 1). ANA: Adaptative Boosting – ANN: Artificial Neural Networks – LR: Logistic Regression – RF: Random Forest – SVM: Support Vector Machine.

Figure 3 shows the bar plots of the predicted and observed infarction volumes one month after stroke for three selected patients. These patients were chosen because they showed discrepancies with SVM. The SVM method underestimated final infarct volume in patient (A), overestimated it in patient (B) and performed similarly to the other methods in patient (C). However, ADA, ANN, LR, and RF performed equally well in all three patients regardless of infarct volume.

**Figure 3.**
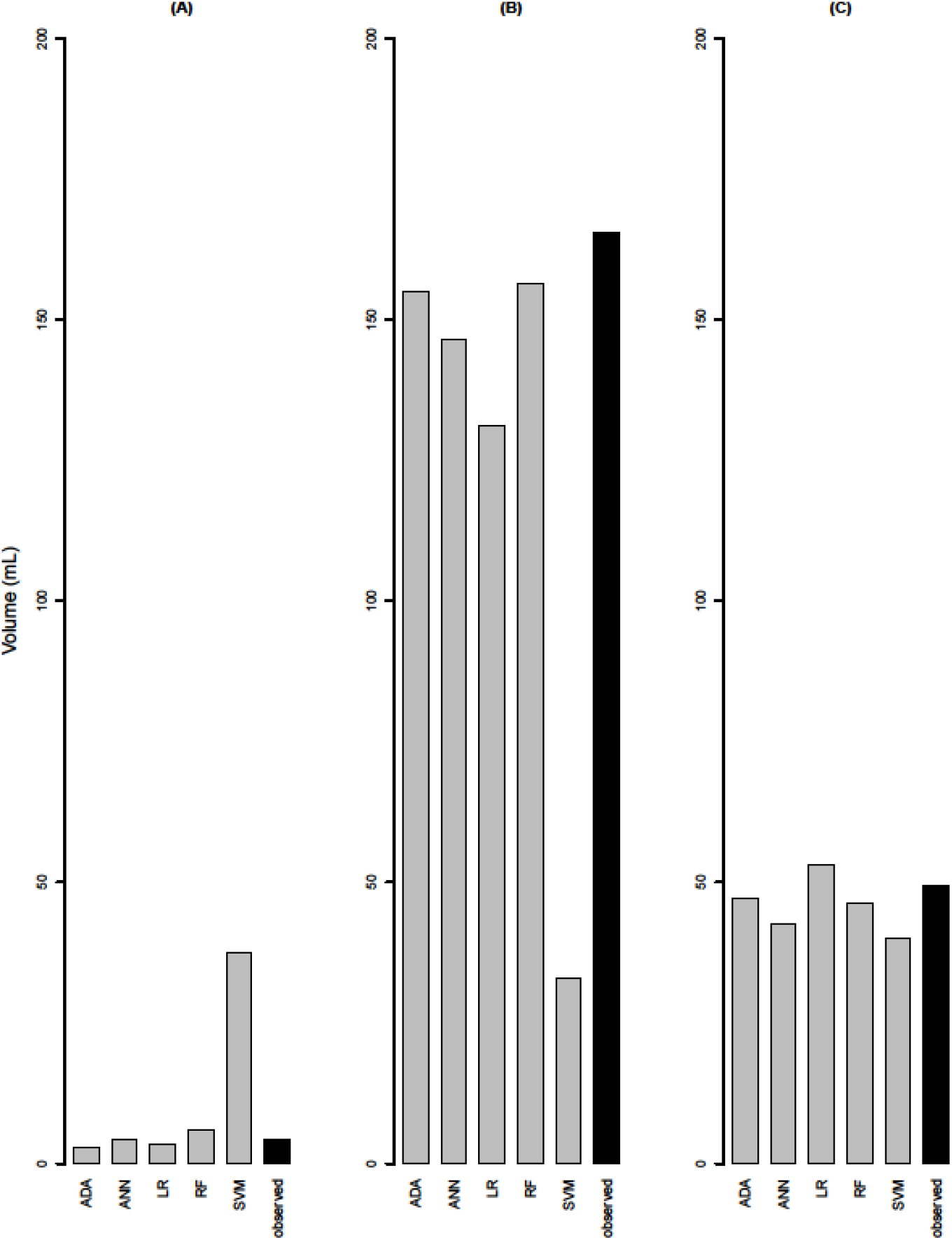
Bar plots of the predicted (gray bars) and observed infarcted volumes (black bars) one month after stroke in three representative patients. ANA: Adaptative Boosting – ANN: Artificial Neural Networks – LR: Logistic Regression – RF: Random Forest – SVM: Support Vector Machine.

## 4. DISCUSSION

To our knowledge, our study is the largest to compare classification methods (especially including four machine learning methods) on imaging data from stroke patients. In this study, five classification methods to identify the brain tissue at risk of infarction were compared using voxel-based multimodal MRI data. Another large study in animals with method comparison was previously reported by Bouts et al.^13^, but was not applied onto human data. Our findings suggest no significant difference in performance of the five classification methods in terms of identification of the tissue at risk of infarction on human imaging data.

Our findings are consistent with previous results obtained based on animal data.^9,13^ However, contrary to our study, a couple studies^10,15^ found that tree-based methods performed better than ANN and GLM. This discrepancy may be due to the use of a different performance criterion, namely Dice coefficient. Dice coefficient considers the true positives and is used to evaluate the accuracy of the predicted infarction volume for a given threshold. However, the comparison metric used in our study *(AUC*_prc_) summarizes the predictive ability over all possible thresholds.

In the present study, the performance of each method was first evaluated by *AUC_prc_* values, which allowed us to summarize the identification ability of the method over all possible thresholds. On the basis of this criterion, the five classification methods performed equally well in identifying tissue at risk of infarction, in agreement with another study that used the same criterion but on animal data.^13^

Regarding the *AUC_roc_* criterion, there was no significant difference in performance between ADA, ANN, LR, and RF methods. However, all performed significantly better than SVM. The latter finding may be due to the fact that SVM is essentially a binary classification method that requires an additional step able to provide the infarction risk. This gives less accurate risk predictions thus lower *AUC_roc_* values.^19^ These results contrast with those obtained on experimental data by Huang et al.^7^ (who showed that SVM outperformed ANN) and with those obtained by Bouts et al.^8^ who showed that all methods performed equally well.

In the present study, *AUC_roc_* values were always higher than *AUC_prc_* values with all methods. This is due to the fact that *AUC_roc_* is not sensitive to the imbalance between healthy and infarcted voxels and overestimate the infarcted volume in studies performed on data with low prevalence.^17^ When we compared the sensitivities and specificities of the methods, the median of sensitivity with each method ranged between 0.4 and 0.5 while the median of specificity was over 0.9. Thus, all methods performed better in identifying healthy tissue than the tissue at risk of infarction.

Most previous studies used mainly the *AUC_prc_*, the *AUC_roc_*, and also the Dice coefficient to compare classification methods on human or experimental ischemic voxel-based data.^8,9,11^ For this reason, we used the more often used criteria in our study: the *AUC_prc_* and the *AUC_roc_*. Regarding the *AUC_prc_*, all previous studies concluded that there were no significant differences between methods. However, RF and SVM performed better than ANN regarding Dice coefficient and *AUC_roc_* criteria in some studies whereas Bouts et al. concluded to similar performances of the methods they used.

The stroke patient data used in this study had a wide range of final infarction volumes. Therefore, the predicted infarcted volumes at one month obtained with different methods were quite different. Despite the heterogeneity of infarcted volumes, ADA, ANN, LR, and RF performed equally and provided homogeneous criteria values whereas SVM gave more heterogeneous values depending on the observed infarction volume.

In terms of computer resources, LR was the fastest method; with one central processing unit (2.7 GHz and 26 Gb RAM), the results were obtained in 30 seconds for all 55 patients whereas SVM required approximately 30 hours. Another advantage of LR vs. other methods is that this method does not need a step of parameter selection to fit the model.

One limitation is that only 55 patients could be included in our study. The performance criteria were calculated using one patient due to leave-one-out cross validation. A bigger data set could let us to use k-fold cross validation, thus to calculate the performance criteria on a test data set including more observation.

In conclusion, the five classification methods performed equally well in terms of identifying the volume of brain tissue at risk of infarction. The ADA, ANN, LR, and RF methods showed equally good performances in term of infarcted volume prediction. The results show that statistical models based on multiparametric MRI can provide valuable prognostic information in acute ischemic stroke, with the potential to guide physicians in their time-critical decision process regarding choice of therapy.

## Acknowledgements

The authors thank Jean Iwaz (Hospices Civils de Lyon) for the revision of the final drafts of the article.

## Author Contribution statement

CT carried out the statistical analyses and drafted the article.

BO organized, checked, and managed the data, helped with the statistical analyses, and commented and reviewed the article.

LO carried out data collection.

D M-B organised the study and commented and reviewed the article.

## Abbreviations

ADA: Adaptative Boosting
ADC: Apparent diffusion coefficient
*AUC_prc_*: Area under the precision-recall curve
*AUC_roc_*: Area under the receiver operating curve
ANN: Artificial Neural Networks
CBV: Cerebral blood volume
CBF: Cerebral blood flow
DWI: Diffusion-weighted imaging
LR: Logistic Regression
MRI: Magnetic Resonance Imaging
MTT: Mean transit time
PWI: Perfusion-weighted imaging
ROC: Receiver operating curve
RF: Random Forest
SVM: Support Vector Machine
TTP: Time to peak
TMAX: Time-to-maximum

