## Supplementary Materials for "Comparison of classification methods for tissue outcome after ischemic stroke"

**APPENDIX**

We explain below the ways the methods used to calculate the infarction risk of a given *n* observation.

The LR estimates the infarction risk of an observation as its probability $\pi_{i}$of being infarcted at one month. Probability $\pi_{i}$ is calculated using MRI parameters $x_{ij}$with the following logit function: $logit$ ( $\pi_{i}$) = $\alpha_{LR}$ + $\sum_{j=1}^{p} \beta_{j,LR}\times x_{ij}$, where $\alpha_{LR}$ is the intercept and $\beta_{j,LR}$ the multiplicative risk factor.

The SVM separates the observations into healthy or infarcted using a linear border. The closest observations to the border in each class are called “support vectors”. The support vectors will help afterwards choosing the best border by maximizing the distance between the border and the support vectors and minimizing the number of misclassified observations. To find a linear border, SVM projects all observations into a higher dimensional space via a kernel function. The SVM optimizes the solution by minimizing $|\left| w | \right|^{2}+ C \sum\xi_{i}$ respecting $y_{i}\left( <\vec{w}.x_{i}>+\alpha\right)\geq1$, where $\vec{w}$ is the normal (perpendicular) vector to the border and C the trade-off constant. The signed distance to the border can be used as the infarction risk. A zero distance represent a voxel that belongs to the border line, a positive distance represents a high risk of infarction, and a negative distance a low risk of infarction.

The ANN is a mathematical representation of natural neural networks. Each network involves several types of layers: an input layer composed of all data $x_{i}$, an output layer that gives the final outcome $y$, and one (or more) hidden layers between the input and the output layer that consist(s) of a set of neurons that process the data and are connected to the other layers. The input layer sends first the information to the next layer with an initial weight and this weight is updated after the response of the network has reached the output layer. The update iterations continue until there is no further change. A weighted sum of the responses of the neurons is then computed as: $sum_{i}=w_{0} +\sum_{j=1}^{p} x_{ij}\times w_{ij}$, where $w_{0}$ is the intercept, $x_{ij}$ the responses of the neurons of the previous layer, and $w_{ij}$ the final updated weights. The risk of infarction is obtained by applying a sigmoid function to this weighted sum: f($x_{i}$) =$\frac{1}{1+exp(-sum_{i})}$.

The RF builds several bootstrap samples from the original data and fits a decision tree to each sample to classify the observations into healthy or infarcted. For each split of a tree classification, the best parameter is chosen within a sample of parameters. This sample is created randomly and its size is fixed a priori (mtry). For a given observation, the risk of infarction is then computed as the percentage of trees that classified this observation as infarcted.

The ADA uses another strategy which corrects sequentially the classification weighting the misclassified observations with a set of decision trees $h_{t}$. The classification of the first tree is performed with the same weight for all observations but the weights of misclassified observations are increased after each decision tree classification. According to the misclassification error of the decision tree and the weights given to the observations, the performance $\alpha_{t}$ of each tree is computed. For a given observation, the infarction risk is the result of the classification of each tree weighted by its performance $\sum_{t=1}^{T} \alpha_{t}\times h_{t}$.

**Settings**

SVM, ANN, RF, and ADA require parameter settings before the fit of the method.

A Gaussian Radial Basis kernel function $K (\vec{x_{1}} ,\vec{x_{2}}) =exp( -\gamma|| \vec{x_{1}} - \vec{x_{2}} ||^2)$ was used for SVM where $\gamma$is a supplementary kernel parameter and $\vec{x_{1}},\vec{x_{2}}$ are observations. To optimize the kernel parameter $\gamma$ as well as the trade-off constant C, a grid-searching approach based on the misclassification error rate was used where C $\in$ [1, 10] was incremented by steps of 1 and $\gamma\in$(0.001, 0.005, 0.01, 0.05, 0.1, 1).

For the ANN, a single hidden layer was fitted and a grid-searching was used to optimize the number of nodes in interval [14, 25] with steps of 1 and a decay parameter $\in$ (0.001, 0.005, 0.01, 0.05, 0.1, 0.5 and 1). The decay parameter can be seen as a regularization parameter that avoids over-fitting.

For the RF, the number of parameters used at each node to perform the classification was set at mrty = 3, the number of trees optimized in the interval [1, 100] by increments of 1, and the performances of the combinations compared using the misclassification error rate.

For the ADA, the optimal number of trees was searched in the interval [1,200] by increments of 1 and the performances of the combinations compared using also the misclassification error rate.

With ADA, RF, ANN, and LR, the threshold that minimizes the difference between the observed and the predicted infarcted volume was set between 0 and 1 with steps of 0.001. With SVM, this threshold was set between -3 and 3 with steps of 0.001.

**Supplementary figures**

Parameter optimization was made using a grid-searching approach that provides the misclassification error rate with ADA, ANN, RF, and SVM algorithms. The parameters which gave the lowest misclassification error rate were chosen.

With ADA, the optimal number of iterations (which is also the number of trees) is searched in the interval [0, 150] with increments of 1. Supplementary figure 1 shows that the misclassification error calculated with ADA is nearly the same when the number of trees is between 100 and 150 and that it reached the minimum at 150 trees when the misclassification error rate was almost $\xi$ = 0.075.

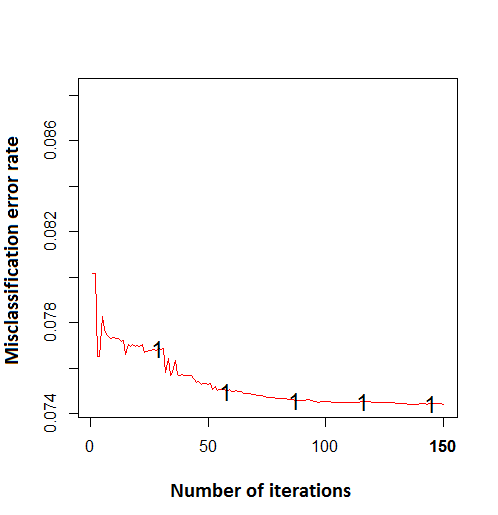

Supplementary figure 1 – Misclassification error rate according to the number of iterations in adaptative boosting analysis.

With RF, the number of randomly chosen variables at each node (mtry) was fixed at 3 as suggested by Breiman and the optimal number of trees (ntree) searched from 1 to 100 with increments of 1 (Freund &, Schapire. *J Japanese Soc Artif Intell* 1999;14:771-780) The misclassification error rate on the Y-axis in Supplementary figure 2 is almost constant for a number of trees ranging between 20 and 100 and the misclassification error rate reached the minimum value at ntree=100. So, the couple of parameters (ntree, mtry) = (100, 3) was used; the misclassification error dropped down to nearly $\xi$ = 0.05.

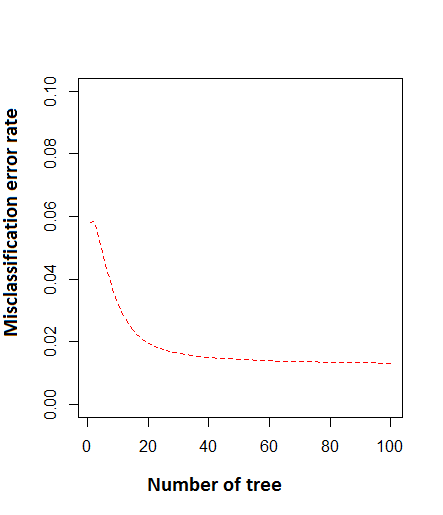

Supplementary figure 2 – Misclassification error rate according to the number of trees in the random forest analysis.

With ANN, the optimization required finding the number of units in the hidden layer. This was searched in the interval [14, 25] with increments of 1 and the optimal decay (learning rate) was searched among values 0.001, 0.01, 0.05, 0.1, 0.5, and 1. Supplementary figure 3 shows the changes of the misclassification error in function of the decay value, the number of units being indicated by different colours. The minimum misclassification error rate ($\xi$= 0.139) was obtained with the parameter couple (size, decay) = (25, 0.1).

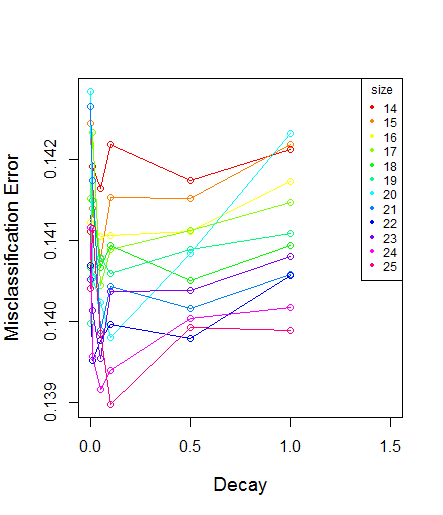

Supplementary figure 3 – Misclassification error rate according to the number of neurons (size) and the decay in artificial neural network analysis.
